# Systematic Mapping of Protein Interactions Underlying IL-2 Secretion in Human T Cells

**DOI:** 10.1101/2025.09.09.675165

**Authors:** Seunghyeon Shin, Frances Rocamora, Nathan E. Lewis

**Affiliations:** Department of Pediatrics, University of California, San Diego, School of Medicine, La Jolla, CA 92093, United States; Department of Bioengineering, University of California, San Diego, La Jolla, CA 92093, United States; Center for Molecular Medicine, Complex Carbohydrate Research Center, and Department of Biochemistry and Molecular Biology, University of Georgia, Athens, GA 30602, United States

**Keywords:** protein-protein interaction, proximity-based labeling, cytokine secretion, secretory pathway, T cell, genetic engineering

## Abstract

Protein secretion plays a crucial role in maintaining immune homeostasis, yet the molecular interactions governing this process remain incompletely understood. While transcriptional and post-transcriptional regulation of protein expression is well characterized, the subcellular interactions between secreted proteins and trafficking machinery are less explored. To address this, we systematically mapped protein–protein interactions (PPIs) involved in the secretion of interleukin-2 (IL-2) from human T cells using proximity-based labeling coupled with mass spectrometry. Our analysis revealed significant enrichment of proteins associated with conventional secretory pathways, including ER-to-Golgi transport, protein folding, and vesicle-mediated trafficking. Functional validation confirmed that several of these proteins are critical for IL-2 secretion, underscoring their direct roles in cytokine processing. In addition, time-resolved profiling of PPIs and transcriptomic changes following T cell stimulation revealed dynamic remodeling of the cytokine secretion machinery, reflecting multilayered regulation at both the protein and gene expression levels. These findings offer a systems-level understanding of IL-2 secretion and identify new molecular components that can be targeted to modulate immune responses. This work provides a framework for dissecting complex secretory processes and has broad implications for therapeutic strategies in immune-related diseases.

## Introduction

Protein secretion is a fundamental process in eukaryotic cells, coordinated by a wide array of secretory machinery proteins distributed across the endoplasmic reticulum (ER), Golgi apparatus, secretory vesicles, endosomes, and the plasma membrane [1]. Most secreted proteins are synthesized with a signal peptide that directs them into the ER-Golgi pathway, also known as the conventional secretory pathway, while some leaderless proteins are secreted via alternative routes that bypass the ER-Golgi complex [2,3]. Proteomic analyses have successfully identified numerous endogenous proteins localized within key secretory compartments, including the rough ER, smooth ER, and Golgi apparatus [4]. Additionally, genome-wide RNAi screens have uncovered sets of genes whose knockdown impairs the secretion of reporter molecules in both Drosophila and human cells, further elucidating the molecular components of the secretory pathway [5,6]. To systematically study protein secretion, genome-scale models of the secretory pathway have been constructed in various cell types, such as yeast, mouse, Chinese hamster ovary (CHO), and human cells, integrating data from genomics, transcriptomics, and proteomics [7–9]. These reconstructed models have enabled the estimation of bioenergetic costs associated with protein secretion and the prediction of target protein productivity, thereby contributing to the optimization of recombinant protein production systems [10,11]. Recent advances in reconstructing the mammalian secretory pathway have enabled systems-level analyses that facilitate the integration of multi-omics datasets and single-cell transcriptomic data [12].

Proximity-based labeling combined with mass spectrometry (MS) has emerged as a robust and versatile approach for identifying protein–protein interactions (PPIs), including transient and weak interactions that are often missed by conventional methods [13]. This technique leverages engineered enzymes, derived from either peroxidases or biotin ligases, to label endogenous proteins in the immediate vicinity of a target protein. In peroxidase-mediated labeling systems, such as horseradish peroxidase (HRP) and APEX, reactive radicals are generated upon enzyme activation, enabling covalent tagging of neighboring proteins within a defined spatial range [14,15]. In contrast, biotin ligase-mediated methods, including BioID and TurboID, utilize a ligase fused to the protein of interest to biotinylate nearby endogenous proteins [16,17]. Each approach offers distinct advantages and limitations. Peroxidase-based methods provide high labeling activity and superior temporal resolution but are often limited to specific cellular compartments and have restricted applicability in in vivo systems due to their dependence on oxidative conditions [18]. On the other hand, biotin ligase-based methods are well-suited for in vivo applications because of their low toxicity, although they typically exhibit lower catalytic efficiency and reduced temporal resolution [18]. The temporal resolution of proximity labeling varies depending on the enzyme used, ranging from minutes to hours, thereby offering either real-time snapshots or cumulative histories of PPIs [19]. By employing short labeling times and targeting engineered enzymes to specific subcellular compartments, both spatial and temporal dynamics of PPIs can be resolved across different time points [20].

Cytokines are small, secreted proteins that are critical in mediating cell-to-cell communication within the immune system, forming a complex and dynamic network among various immune cell types [21]. Over a hundred cytokines have been identified in the human immune system, and their expression is tightly regulated by the coordinated action of multiple transcription factors [22]. While the transcriptional and post-transcriptional regulation of cytokines has been extensively studied, the post-translational regulation, particularly the mechanisms underlying their secretion, remains poorly understood [23–26]. Cytokines can be secreted through both canonical and non-canonical pathways, with the former involving trafficking through the ER-Golgi complex and the latter utilizing alternative secretory compartments [27]. Cytokines that contain signal peptides and glycosylation sites are typically directed into the ER-Golgi pathway, whereas those lacking signal peptides are secreted via unconventional routes [28]. Several families of trafficking machinery proteins, including soluble N-ethylmaleimide-sensitive-factor attachment protein receptor (SNARE) proteins, Rho and Rab GTPases, and Golgins, are involved in mediating cytokine secretion [28–30]. However, the study of cytokine secretion mechanisms has largely relied on prior knowledge and has often focused on individual components of the secretory machinery. With the advent of high-throughput techniques such as proximity-based labeling, it is now possible to systematically map the PPIs of specific target proteins within cells, providing a more comprehensive understanding of the cellular machinery involved in cytokine secretion.

Interleukin-2 (IL-2) is a key regulator of T cell-mediated immune responses. Its gene expression is tightly controlled by a network of transcription factors, including AP-1, NFAT, and NF-κB, which are activated downstream of T cell receptor (TCR)/CD3 and CD28 co-stimulatory signaling pathways [31,32]. IL-2 exerts its effects by binding to the IL-2 receptor (IL-2R), which exists in three configurations: the low-affinity IL-2Rα monomer, the intermediate-affinity IL-2Rβ/γ heterodimer, and the high-affinity IL-2Rα/β/γ heterotrimer [33]. While IL-2Rβ and IL-2Rγ are constitutively expressed in T cells, the expression of IL-2Rα is inducible upon TCR-mediated stimulation [34]. IL-2 signaling triggers a wide range of immune responses across multiple cell types, including CD4+ and CD8+ T cells, regulatory T (Treg) cells, B cells, and natural killer (NK) cells [35]. Depending on its concentration and context, IL-2 can promote the expansion and activation of either effector T cells (at transient high levels) or Treg cells (at chronic low levels) [36]. Due to its pleiotropic effects on various immune cells expressing different IL-2 receptor forms, IL-2 has become a therapeutic target for both cancer and autoimmune diseases [37]. It is also one of the few cytokines approved by the FDA for cancer immunotherapy, used to treat conditions such as renal carcinoma and metastatic melanoma [38]. However, despite its therapeutic potential, the clinical use of IL-2 has been limited by its high toxicity and broad activity, which have prompted the development of engineered IL-2-based therapies, including IL-2 muteins, mimetics, PEGylated IL-2, IL-2/antibody complexes, and IL-2 fusion proteins, to enhance specificity and reduce side effects [39]. In this context, a detailed understanding of the PPIs involved in IL-2 secretion may offer valuable insights into the mechanisms governing cytokine trafficking and identify novel targets for engineering IL-2 variants that better balance immune stimulation and suppression.

In this study, we employed a proximity-based labeling approach to identify PPIs involved in the secretion of IL-2 from human immune cells. Analysis of the biotinylated proteins in close proximity to IL-2 revealed strong enrichment in components of the conventional secretory pathway, particularly within the ER and Golgi apparatus. Functional perturbation of genes corresponding to these enriched PPIs led to a marked decrease in IL-2 secretion, confirming their roles in the secretory process. In addition, time-resolved PPI profiling following T cell stimulation indicated that many proteins are recurrently engaged in IL-2 secretion across multiple time points. Finally, transcriptomic analyses highlighted the multilayered regulatory mechanisms underlying IL-2 production and secretion, reflecting the dynamic and tightly controlled nature of cytokine expression in the human immune system.

## Experimexntal Section

### Cell Culture

Jurkat T cells (Clone E6-1, ATCC) were cultured in RPMI 1640 (ATCC) supplemented with 10% fetal bovine serum (Thermo Fisher Scientific) and penicillin-streptomycin (Thermo Fisher Scientific). Cells were passed every 2 to 3 days, seeded at a cell density of 0.3 to 0.5 x 10^6^ cells/mL.

### Cell Stimulation

Jurkat T cells were stimulated using the culture medium supplemented with PMA (Sigma-Aldrich), ionomycin (Thermo Fisher Scientific), PHA (Sigma-Aldrich), and ImmunoCult™ Human CD3/CD28 T Cell Activator (STEMCELL Technologies). The stocks of PMA and ionomycin were diluted with DMSO, and the stock of PHA was diluted with distilled water, aliquoted and stored at -20 ⁰C.

### Western Blot

Cells were washed with PBS twice and lysed using RIPA lysis buffer (Thermo Fisher Scientific) supplemented with protease inhibitor cocktail (Sigma-Aldrich) according to the manufacturer’s instructions. Protein concentrations were measured using a BCA protein assay kit (Thermo Fisher Scientific) according to the manufacturer’s instructions. Samples were boiled with a sample buffer (Bio-Rad) and beta-mercaptoethanol (Bio-Rad) at 95 ⁰C, 5 min. Samples were loaded in precast gels, run, and transferred to nitrocellulose membranes using the Trans-Blot Turbo system (Bio-Rad) according to the manufacturer’s instructions. The membranes were blocked in a blocking buffer (LICORbio) for 1 hour. Primary and secondary antibodies were diluted in antibody diluent (LICORbio) and incubated with membranes, followed by washing in 0.1% TBST. For the visualization of proteins, SuperSignal™ West Femto Substrate (Thermo Fisher Scientific) was used, and images were analyzed using G:BOX Chemi XRQ gel doc system (Syngene). For staining of biotinylated proteins, 5 μg of protein were loaded and transferred into nitrocellulose membranes as previously described. The membranes were blocked in 3% BSA with 0.1% TBST for 1 hour. Streptavidin-HRP (Cell Signaling Technology), diluted in blocking solution at 1:2000, was incubated with membranes, followed by wash in 0.1% TBST.

### Proximity-Based Labeling

Biotinylation by antibody-guided recognition (BAR) method was used to label endogenous proteins in proximity with IL-2 proteins [40]. Stimulated or not-stimulated Jurkat T cells were harvested, and cells were washed with 0.1% PBST, fixed with 4% PFA (Thermo Fisher Scientific), and permeabilized with 0.4% PBST. Endogenous HRP activity was blocked by treating 0.4% H_2_O_2_ (Thermo Fisher Scientific) for 10 min, and cells were blocked with a blocking buffer, which is 5% goat serum (Gibco) with 1% BSA in 0.1 PBST, for 1 hour. The primary antibody targeting IL-2 (ABclonal) was diluted in a blocking buffer at 1:500, and incubated with samples overnight at 4 ⁰C. The secondary antibody conjugated with HRP (Abcam) was diluted in a blocking buffer at 1:1000, and incubated with samples for 1 hour. Cells were incubated with biotin-tyramide (Akoya Biosciences) and amplification diluent (Akoya Biosciences), followed by reaction stop with 0.5 M Na ascorbate (Thermo Fisher Scientific). Samples were boiled with SDS (Thermo Fisher Scientific)and sodium deoxycholate (Thermo Fisher Scientific) at 99 ⁰C for 1 hour.

### Fluorescence Microscopy

The co-localization of biotinylated proteins and IL-2 proteins was observed using Leica SP8 Confocal with Lightning Deconvolution (Leica Microsystems). Streptavidin-DyLight 594 (Thermo Fisher Scientific) and goat anti-rabbit DyLight 650 (Abcam) were diluted in blocking buffer, which is 5% goat serum (Gibco) with 1% BSA, at 1:1000 and 1:250, respectively, and incubated with biotinylated samples from BAR experiments for 0.5 hour. After washing three times with 0.1% PBST, cells were resuspended with mounting media (Vector Laboratories) and mounted on the slide.

### Affinity Purification

Affinity purification was performed on the Bravo AssayMap platform (Agilent) using AssayMap streptavidin cartridges (Agilent). Cartridges were initially primed with 50 mM ammonium bicarbonate, followed by the slow loading of protein samples onto the streptavidin matrix. To eliminate background contaminants, the cartridges were washed with 8 M urea in 50 mM ammonium bicarbonate. Subsequently, they were rinsed with Rapid Digestion Buffer (Promega), and on-cartridge digestion was carried out using mass spectrometry-grade Trypsin/Lys-C Rapid Digestion Enzyme (Promega) at 70 °C for 1 hour. The resulting peptides were desalted on the Bravo platform using AssayMap C18 cartridges and dried in a SpeedVac concentrator.

### Mass Spectrometry Analysis

Prior to LC-MS/MS analysis, dried biotin-enriched peptides were reconstituted in 2% acetonitrile (ACN) and 0.1% formic acid (FA), and their concentrations were determined using a NanoDrop spectrophotometer (Thermo Fisher Scientific). Peptide samples were analyzed using a Proxeon EASY-nanoLC system (Thermo Fisher Scientific) coupled to an Orbitrap Fusion Lumos Tribrid mass spectrometer (Thermo Fisher Scientific). Separation was performed on a C18 Aurora analytical column (75 µm × 250 mm, 1.6 µm particle size; IonOpticks) at a flow rate of 300 nL/min, using a 75-minute gradient as follows: 2–6% buffer B in 1 min, 6–23% B over 45 min, 23–34% B over 28 min, and 34–48% B in 1 min (buffer A: 0.1% FA; buffer B: 80% ACN with 0.1% FA). The mass spectrometer operated in positive ion data-dependent acquisition (DDA) mode. MS1 spectra were acquired in the Orbitrap over a mass-to-charge (m/z) range of 375–1500 at a resolution of 60,000, with an automatic gain control (AGC) target of 4 × 10⁵ and a maximum injection time of 50 ms. The instrument was set to top-speed mode with a 1-second cycle time for MS1 and MS/MS scans. The most intense precursors (charge states +2 to +7) were selected for fragmentation using higher-energy collisional dissociation (HCD) at 30% normalized collision energy, with precursor isolation performed in the quadrupole using a 0.7 m/z window. MS/MS spectra were acquired in the ion trap in rapid scan mode with an AGC target of 1 × 10⁴ and a maximum injection time of 35 ms. Dynamic exclusion was enabled for 20 seconds with a 10 ppm mass tolerance around the precursor m/z.

### MS Data Analysis

All mass spectra were analyzed using MaxQuant software (version 1.6.11.0) [41]. MS/MS spectra were searched against the Homo sapiens UniProt protein database (downloaded in July 2023) supplemented with GPM cRAP sequences, which represent common protein contaminants. The precursor mass tolerance was set to 20 ppm for the initial search (used for mass recalibration) and 4.5 ppm for the main search. Product ions were searched with a mass tolerance of 0.5 Da. The maximum precursor ion charge state considered was 7. Trypsin was specified as the digestion enzyme with specific cleavage rules, allowing up to two missed cleavages. A target-decoy approach was used to control the false discovery rate (FDR), which was set at 1%. Differential analysis of biotinylated proteins between samples was conducted in Perseus software [42], based on label-free quantification (LFQ) intensity values.

### siRNA-mediated Knockdown

Transfection of Jurkat T cells was performed using Amaxa Nucleofector II (Lonza) and Cell Line Nucleofector V kit (Lonza). Silencer Select siRNAs (Thermo Fisher Scientific) for each target gene were transfected into cells using the X-005 program, according to the manufacturer’s instructions. Cells were washed twice with PBS, and 5 x 10^6^ cells per cuvette were subjected to nucleofection with 500 nM siRNA.

### qPCR

Total RNA was extracted using the RNeasy Plus Mini Kit (Qiagen), according to the manufacturer’s instructions. RNA concentrations were measured using Nanodrop (Thermo Fisher Scientific), and cDNA was synthesized from 1 μg of total RNA using SuperScript™ II Reverse Transcriptase (Thermo Fisher Scientific), according to the manufacturer’s instructions. RT-qPCR was performed using iTaq Universal SYBR Green Supermix (Bio-Rad) and CFX96™ Real-Time System (Bio-Rad) with the following amplification program: 95 ⁰C for 30 sec, 40x: 95 ⁰C for 5 sec, 60 ⁰C for 1 min. Relative expression of mRNA was measured using the ΔΔCt method and normalized with human ACTB gene expression.

### RNA Sequencing and Data Analysis

Total RNA quality was evaluated using the Agilent TapeStation 4200, and only samples with an RNA Integrity Number (RIN) above 8.0 were selected for library preparation. RNA-seq libraries were generated using the Illumina Stranded mRNA Prep kit (Illumina), following the manufacturer’s protocol. Prepared libraries were multiplexed and sequenced on an Illumina NovaSeq X Plus platform using 150 bp paired-end (PE150) reads, targeting a sequencing depth of approximately 25 million reads per sample. Demultiplexing was performed with Illumina’s bcl2fastq Conversion Software (Illumina). Raw sequencing reads in FASTQ format were first assessed for quality using FastQC [43], followed by adapter and barcode trimming using TrimGalore [44]. Transcript quantification was performed by pseudoalignment with Kallisto [45], enabling fast and accurate transcript-level abundance estimation from the processed reads.

### Enzyme-Linked Immunosorbent Assay (ELISA)

Human IL-2 DuoSet ELISA kit (R&D Systems) was used to measure the secreted IL-2 proteins when cells were stimulated, according to the manufacturer’s instructions. Each time point of post-stimulation, samples were prepared by centrifugation of cell culture media, and supernatant was stored at -80 ⁰C for ELISA.

### Statistical analysis

All statistical analyses were conducted using GraphPad Prism (version 10.4.1), unless specified otherwise. For comparisons between two independent groups, statistical significance was evaluated using an unpaired, two-tailed Student’s t-test. For comparisons involving more than two groups, one-way analysis of variance (ANOVA) followed by Tukey’s post hoc test was applied. Data are reported as mean ± standard deviation (SD), unless otherwise noted. A p-value of less than 0.05 was considered statistically significant.

## Results

### Validation of BAR method for IL-2 secretion

To identify the PPIs during IL-2 secretion, the BAR method, which is a proximity-based labeling method using antibody-guided biotinylation, was used in Jurkat T cells. BAR method exploits biotinylation mediated by antibody-conjugated HRP that can bind to target proteins, resulting in the biotinylation of endogenous proteins in proximity with target proteins [40]. Then, biotinylated proteins are captured and identified by MS analysis, providing interactome data of target proteins. The concentration of secreted IL-2 was measured using ELISA when Jurkat T cells were stimulated, and IL-2 secretion level was saturated after 24 hours from stimulation (Figure S1). Intracellular IL-2 was monitored to determine the time point for PPI detection, showing the increase of IL-2 until the 6 hours of stimulation then slowly decreasing (Figure S1). For the validation of the BAR method, the distribution patterns of IL-2 and biotinylated proteins were observed using fluorescence microscopy. Intracellular IL-2 and biotinylated proteins were only observed in stimulated cells, and those proteins were highly colocalized in specific cellular compartments, indicating the selective biotinylation induced by IL-2 recognition (Figure 1a). IL-2 proteins are secreted from T cells into the immunological synapse through the secretory compartments after the stimulation by the engagement with antigen-presenting cells [46]. Here, IL-2 proteins also exhibited a certain directionality of secretion unlike other recombinant proteins [47]. Finally, biotinylation of endogenous proteins were observed using streptavidin blotting, and the sample with 6 hours of stimulation that incubated with biotin reagent showed a significant increase in biotinylation compared to control (Figure 1b).

**Figure 1.**
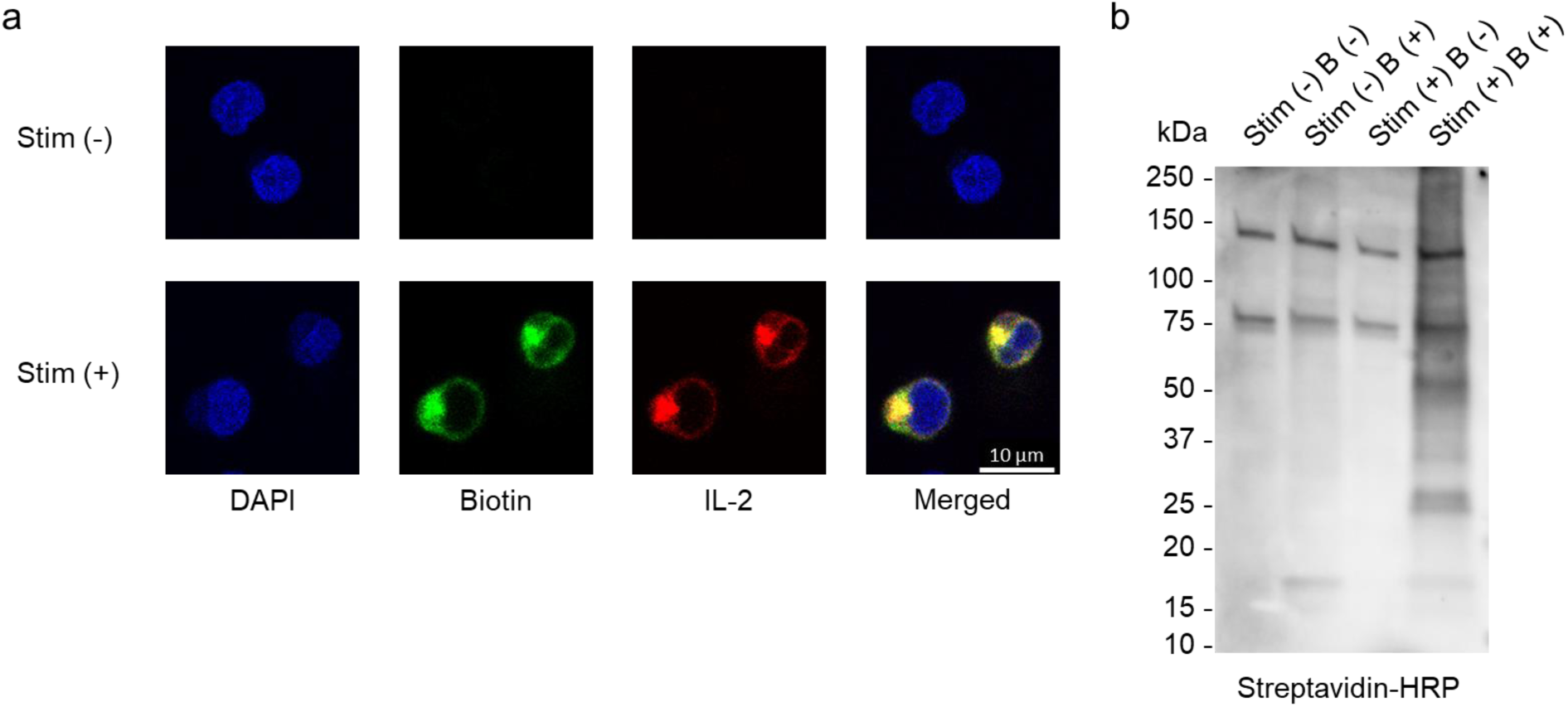
Validation of proximity-based labeling method for IL-2 secretion in human T cells. (a) Co-localization of IL-2 with biotinylated proteins labeled using the HRP-mediated proximity labeling method. Intracellular biotinylated proteins were visualized with streptavidin-DyLight 594 (green), and IL-2 was stained with DyLight 650 (red). (b) Detection of endogenous protein biotinylation by streptavidin blotting. Biotinylation profiles were assessed in stimulated (Stim+) and non-stimulated (Stim−) samples, in the presence (B+) or absence (B−) of biotin reagent.

### Identification of IL-2 PPIs using the BAR method

Given that the level of intracellular IL-2 proteins is highest after 6 hours of stimulation, we decided to collect the MS samples at this time point in triplicate parallel with the control sample without stimulation (Figure 2a). Cells were fixed, permeabilized, blocked, and treated with antibodies binding to IL-2. HRP conjugated with secondary antibodies mediates chemical reaction converting biotin tyramide into highly reactive radicals that can be attached to the endogenous proteins in proximity with IL-2. Biotinylated proteins were captured using streptavidin cartridges, and peptides were obtained from on-cartridge digestion, subjected to LC-MS/MS analysis (Figure 2a). First, several endogenous proteins, which are known to be biotinylated, were detected in both control and IL-2 secretion samples, including acetyl-CoA carboxylase 1 and 2 (ACACA and ACACB), methylcrotonoyl-CoA carboxylase alpha and beta (MCCC1 and MCCC2), propionyl-CoA carboxylase alpha and beta (PCCA and PCCB), and pyruvate carboxylase (PC). Then, the differences of biotinylated proteins between control and IL-2 secretion samples were analyzed using Perseus [42], based on the LFQ intensity value of each sample. After the imputation of MS data by replacing missing values from a normal distribution, a total of 366 proteins were enriched in IL-2 secretion samples compared to the control samples (p-value < 0.01 and Log_2_(Fold Change) > 1), including proteins related to protein transport from ER to Golgi and protein folding (Figure 2b).

**Figure 2.**
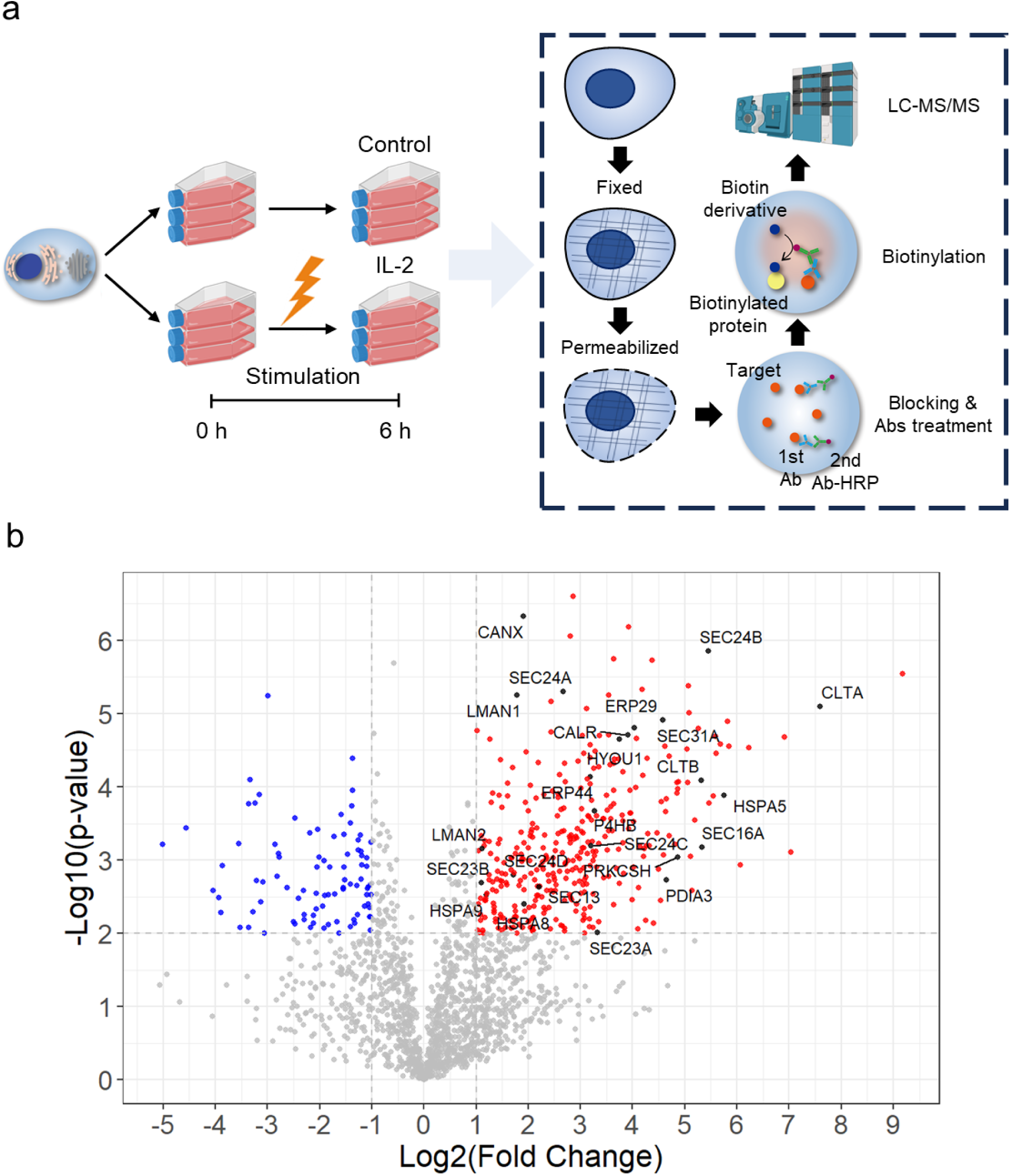
Profiling of enriched protein–protein interactions putatively involved in IL-2 secretion. (a) Schematic overview of the sample preparation workflow for mass spectrometry and the biotin labeling procedure. Created with <u>BioRender.com</u>. (b) Volcano plot of IL-2-interacting proteins. Proteins significantly enriched (p-value < 0.01 and Log₂(Fold Change) > 1) are shown in red, while depleted proteins are shown in blue. Proteins associated with ER-to-Golgi transport and protein folding are annotated.

### Analysis of enriched secretory pathways

Gene ontology (GO) annotation analysis using a total of 366 proteins, which were detected in IL-2 secretion samples, revealed that several biological processes were significantly enriched, including protein transport, cytoskeleton organization, multivesicular body assembly, and stress granule assembly (Figure 3a). Vesicle-mediated transport was also significantly enriched, such as the COPII-coated vesicle, suggesting the directionality of secretion from ER to Golgi. GO terms of cellular components were enriched in secretory pathways, including ER, Golgi, plasma membrane, cytoskeleton, and exosome (Figure 3a). Enriched secretory pathways are highly related to the canonical secretory pathway, which is known to be exploited by IL-2 secretion, passing through the ER and Golgi complex to plasma membrane [27]. To enrich the secretory machinery genes governing IL-2 secretion, we filtered out genes not related to the secretory pathway using the list of proteins previously identified (n=575) in human cells [9]. As a result, a total of 47 proteins were obtained and subjected to PPI network analysis using the STRING database [48]. The physical subnetwork of PPIs was highly enriched (PPI enrichment p-value < 1.0e-16), indicating that secretory machinery proteins biotinylated by the BAR method are localized in proximity with each other, contributing to the IL-2 secretion (Figure 3b). Then, the Markov clustering (MCL) algorithm (inflation parameter: 1.7) was used to identify the function of each protein complex, including protein folding, COPII-coated vesicle cargo loading, SNARE interactions in vesicular transport, COPI-mediated anterograde transport, clathrin coat of trans-Golgi network vesicle, and ubiquitin-dependent ER-associated protein degradation (ERAD) pathway (Figure 3b).

**Figure 3.**
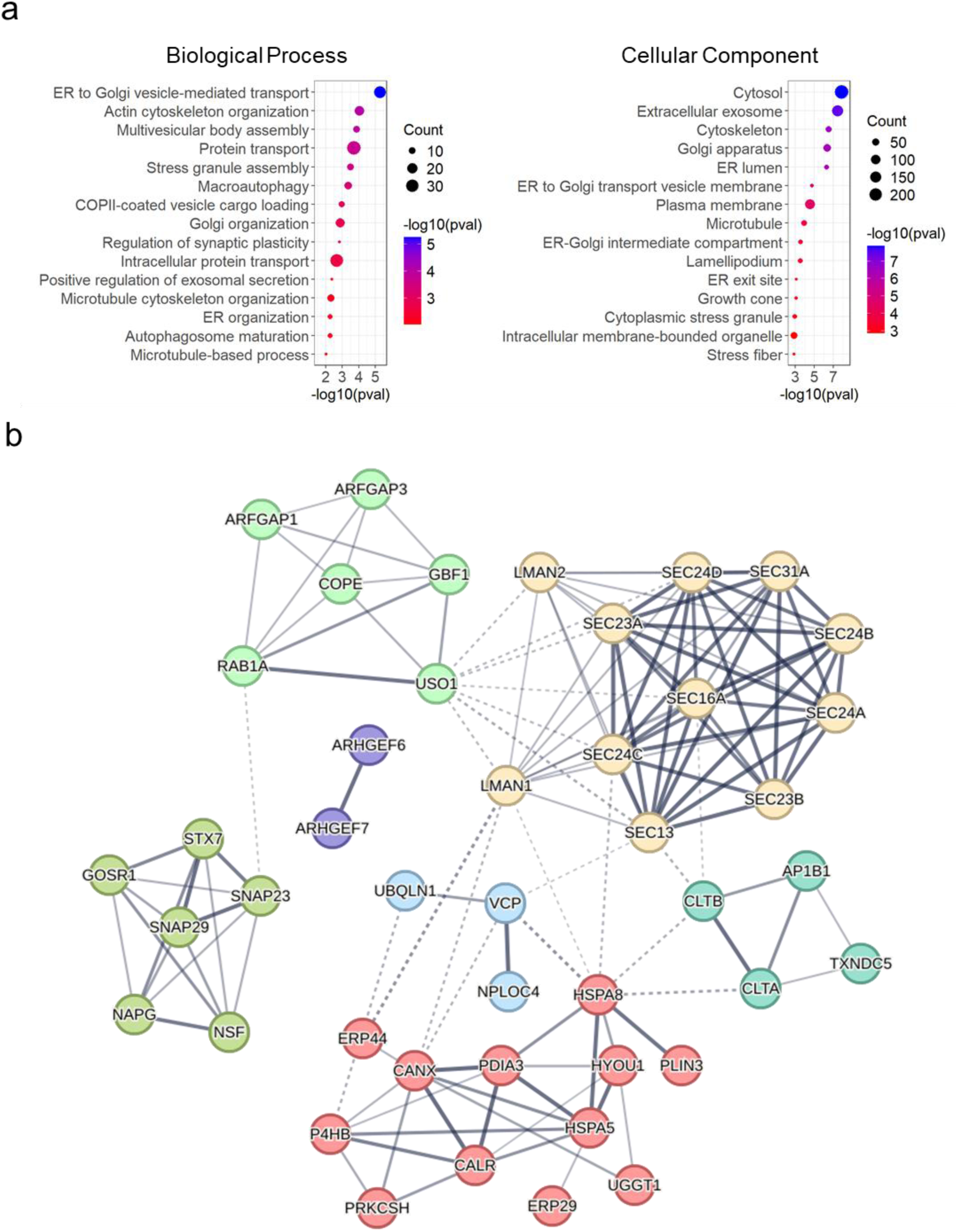
Enriched biological pathways of IL-2–interacting proteins. (a) Gene ontology (GO) analysis of enriched protein–protein interactions categorized by biological process and cellular component. All proteins detected using the BAR method (n=1,895) were used as the background set, and the top 15 GO terms were ranked by p-value. (b) Physical interaction subnetwork of biotinylated proteins generated using the STRING database. Among 366 biotinylated proteins, 47 were identified by filtering against a previously curated human secretory pathway database (n=575). A total of 44 proteins were clustered using the Markov clustering (MCL) algorithm (inflation parameter=1.7) as implemented in STRING.

### Validation of target genes using siRNA-mediated gene knockdown

To evaluate the effects of identified genes on IL-2 secretion, siRNA was used to decrease the expression of each gene (Figure 4a). First, cells were seeded separately for transfection 24 hours in advance, then siRNA targeting each gene was treated to decrease the mRNA expression of target genes in cells. The level of mRNA expression was measured after 48 hours of incubation, which is the time point exhibiting higher siRNA efficiency (Figure S2). Cells were stimulated to secrete IL-2 after another 48 hours incubation, providing enough time for the degradation of previously-expressed proteins. Finally, ELISA samples were prepared after 24 hours of stimulation to assess the effect of gene knockdown. We tested the effect of gene knockdown on IL-2 secretion using a total of 42 genes from 6 clusters compared to the non-targeting siRNA (Figure 4b). In cluster 1, several proteins related to unfolded protein binding and protein disulfide isomerase (PDI) activity exhibited significant reduction of IL-2, such as CALR, CANX, ERP29, ERP44, HSPA5, HSPA8, HYOU1, P4HB, PDIA3, and UGGT1 (Figure 4c). In cluster 2, both LMAN1 and LMAN2, which are lectin proteins binding to glycoproteins and mediating trafficking of them, showed significant reduction and two members of the SEC24 protein family, SEC24A and SEC24D, showed increased IL-2 concentration (Figure 4d). In cluster 3, among the six proteins involved in membrane fusion, three proteins, GOSR1, NAPG, and NSF, which mediate intra-Golgi vesicle-mediated transport, decreased IL-2 secretion (Figure 4e). In Cluster 4, two proteins involved in COPI-coated vesicle budding, COPE and GBF1, increased the concentration of secreted IL-2, while USO1, which is essential for vesicle tethering, decreased IL-2 levels (Figure 4f). In cluster 5, both clathrin light chain proteins, CLTA and CLTB showed reduced IL-2 secretion (Figure 4g). In cluster 6, NPLOC4 and VCP, which form ternary complexes with UFD1 and play an important role in ubiquitin-mediated protein degradation, exhibited decreased IL-2 secretion (Figure 4h). Overall, 24 out of 42 genes showed significant changes in IL-2 secretion from T cells, and most of them decreased the IL-2 secretion and only 4 genes (SEC24A, SEC24D, COPE, and GBF1) increased the IL-2 secretion. Notably, genes related to protein folding exhibited a higher reduction of IL-2 secretion, highlighting the major role of molecular chaperones in cytokine secretion (Figure 4c). Among the genes within the protein folding cluster, HSPA8 knockdown resulted in the most significant reduction in IL-2 secretion. Upon stimulation, HSPA8 proteins were found to co-localize with IL-2, suggesting that HSPA8 plays a role in IL-2 secretion, likely through protein-protein interactions (Figure S3).

**Figure 4.**
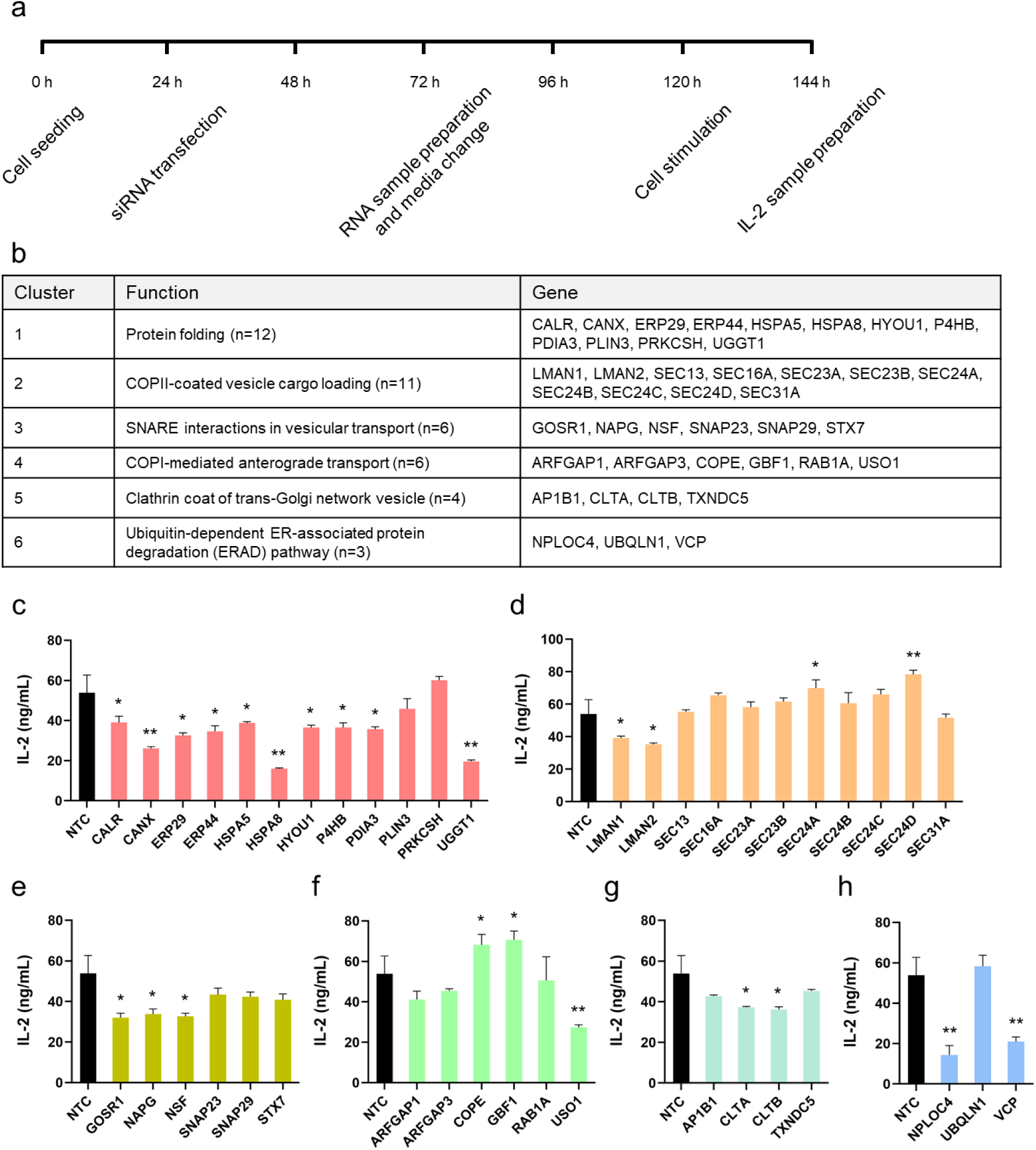
Effects of gene knockdown on IL-2 secretion. (a) Overview of the siRNA-mediated gene knockdown workflow. (b) Gene lists from each cluster associated with secretory pathways and significantly enriched in IL-2 interactions. (c–h) Effects of individual gene knockdown on IL-2 secretion compared to non-targeting siRNA control (NTC) for clusters 1 (c), 2 (d), 3 (e), 4 (f), 5 (g), and 6 (h).

### Temporal analysis of PPIs during IL-2 secretion

To understand time-dependent changes of PPIs after T cell stimulation, the BAR method was performed in four samples, including control without stimulation (T0), 4 h (T1), 6 h (T2), and 8 h (T3) after stimulation (Figure 5a). Different time points were selected based on the intracellular IL-2 protein expression (Figure S1). Assessment of similarity and variability of PPIs between samples showed separation between control and stimulated samples, and T1 (early phase) exhibited higher clustering compared to T2 (middle phase) and T3 (late phase), which are very similar to each other in PCA plot and heatmap analysis (Figure S4). Among the enriched PPIs in each sample, 56 PPIs were consistently enriched at all three points, 40 and 32 PPIs were only enriched at T2 or T3, respectively (p-value < 0.01 and Log_2_(Fold Change) > 1) (Figure 5b). Notably, 56 proteins exhibit a highly significant PPI enrichment p-value (< 1.0e-16), and enriched biological processes, including protein transport and folding (Figure S5). Additionally, 40 and 32 proteins, detected exclusively at T2 or T3, respectively, show lower enrichment p-values compared to proteins consistently enriched across all time points (Figure S5). This suggests that protein interactions related to IL-2 transport and folding remain consistent throughout the early, middle, and late phases of secretion. To investigate the non-secretory pathway interactions, we filtered out genes related to the secretory pathway database [9] from the list of genes showing consistent interactions at all time points, resulting in 45 genes (Figure 5c). Enriched proteins are mainly observed in other cellular compartments, such as extracellular exosome and early endosome, and a total of 15 proteins (PTPN1, TFRC, HDLBP, PEBP1, ANXA11, ZC3HAV1, CORO1A, TUFM, ATXN2L, CSDE1, CAPRIN1, GOLGB1, STRAP, UBAP2L, EIF4B) has molecular function as an RNA binding protein (Figure 5c). Among the RNA binding proteins, 5 proteins (ANXA11, ATXN2L, CAPRIN1, CSDE1, and UBAP2L) are involved in stress granule assembly [49–53]. Cytokine production is regulated in a post-transcriptional manner by RNA binding proteins and the formation of stress granules, because of adenine and uridine-rich elements (AREs) in cytokine mRNA [54,55]. Therefore, we decided to focus on the effects of RNA binding proteins on IL-2 secretion when their expression is inhibited. To assess the effects of RNA binding proteins on IL-2 secretion, cells were seeded and treated with siRNA for each target gene, followed by cell stimulation. After stimulation, samples were prepared at 8 and 24 h, which are the exponential and saturated phase for IL-2 secretion, respectively (Figure S1). As a result, the knockdown of ANXA11, ATXN2L, and GOLGB1 genes showed reduced IL-2 secretion at 8 h after stimulation, and the knockdown of ANXA11, ATXN2L, and UBAP2L genes showed reduced IL-2 secretion at 24 h after stimulation (Figure 5d).

**Figure 5.**
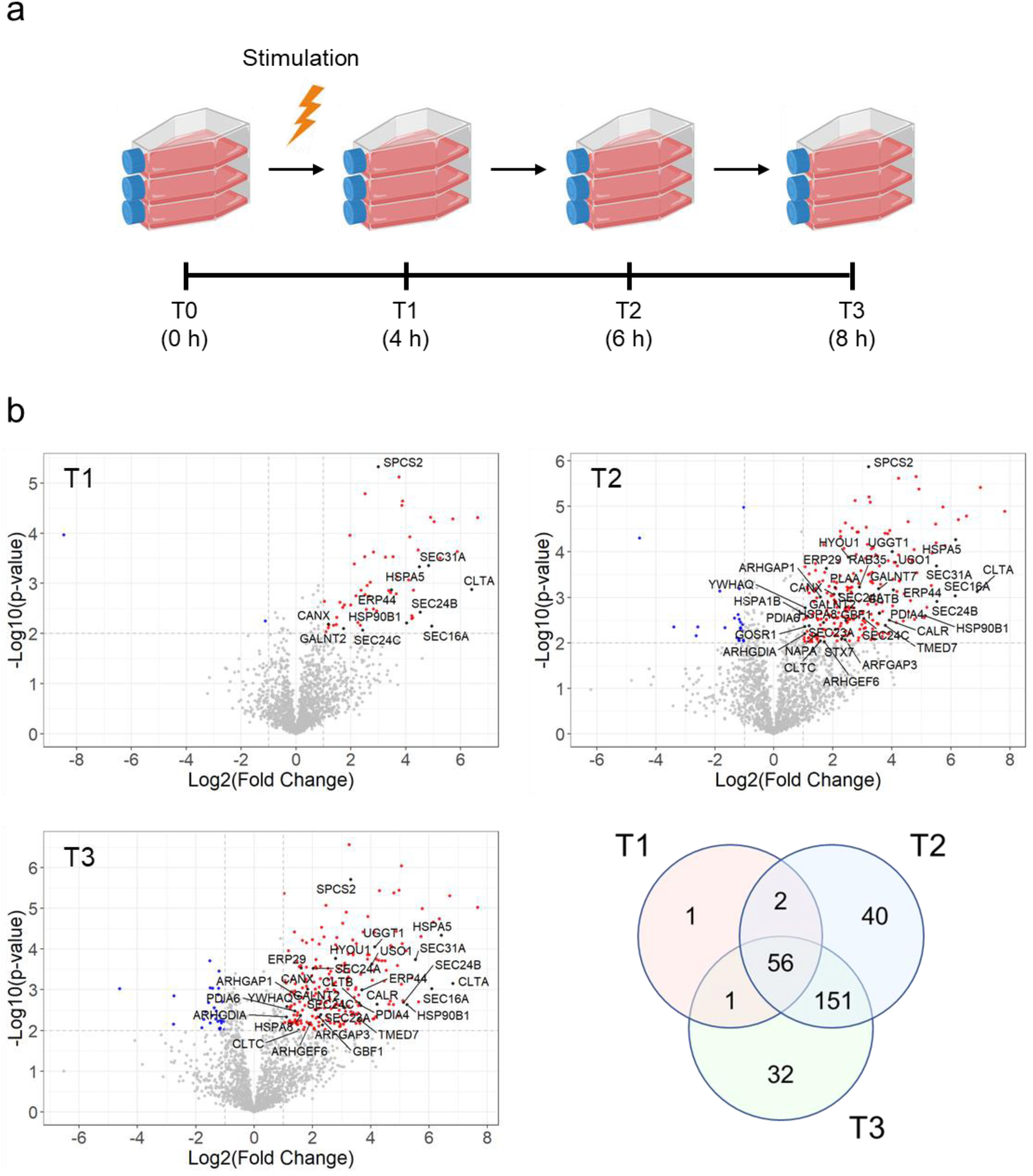
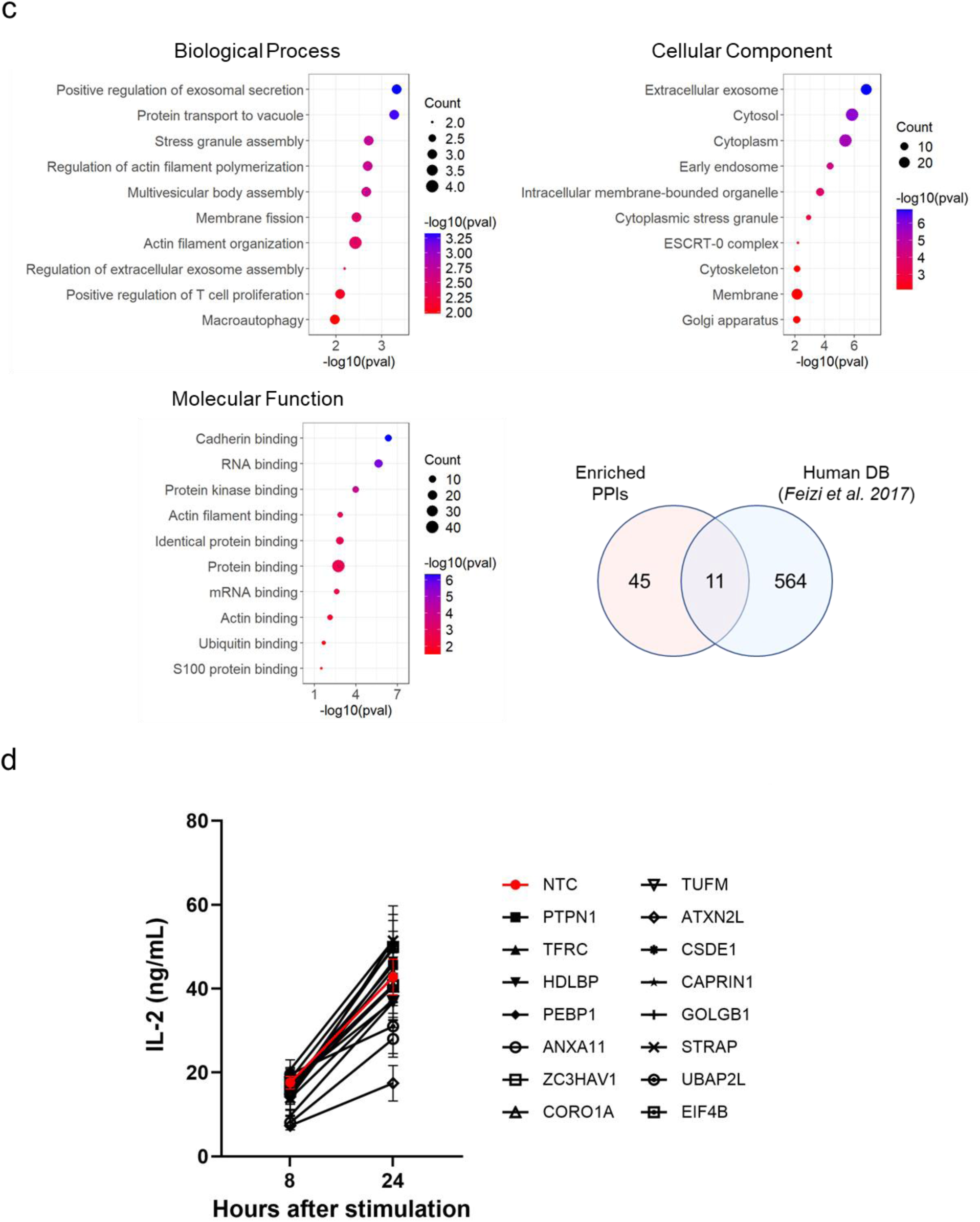
Time-resolved analysis of enriched protein–protein interactions. (a) Schematic overview of the sample preparation workflow. (b) Enrichment of IL-2–interacting proteins at each time point (T1, T2, and T3) compared to the control (T0). Proteins significantly enriched (p < 0.01 and Log₂(Fold Change) > 1) are shown in red; depleted proteins are shown in blue. Proteins annotated in the human secretory pathway database (n=575) are labeled. (c) Gene ontology analysis of 45 proteins consistently enriched at all three time points (T1, T2, and T3) and not associated with the human secretory pathway. (d) Effects of siRNA-mediated knockdown of RNA-binding proteins enriched at all three time points (T1, T2, and T3) on IL-2 secretion at 8 or 24 hours after stimulation.

### Multilayered regulation of IL-2 secretion

Cytokine secretion is temporally and spatially regulated by the expression of multiple genes when immune cells are stimulated [56]. To understand this process systematically, RNA sequencing was conducted in four samples, including control without stimulation (T0), 4 h (T1), 6 h (T2), and 8 h (T3) after stimulation, same as PPIs data set (Figure 5a). Similar to the proteomic analysis, there is a separation between control and stimulated samples, and higher clustering between T2 and T3 (Figure S6). Across all time points, 858 genes were upregulated (p-value < 0.05 and Log_2_(Fold Change) > 0) and 764 genes were downregulated (p-value < 0.05 and Log_2_(Fold Change) > 0) in stimulated samples compared to control samples (Figure 6a). To assess the effects of differential RNA expressions on PPIs, we generated the list of proteins which are significantly enriched in PPIs with increased or decreased RNA expression levels. As a result, 81 genes were identified with increased RNA expression levels and enriched PPIs with IL-2 (Figure 6b). GO annotation analysis revealed that these genes were significantly associated with biological pathways such as ER-to-Golgi protein transport and protein folding in the ER (Figure 6c). Similarly, 60 genes were identified with decreased RNA expression levels and enriched PPIs with IL-2 (Figure 6b), which were significantly associated with pathways related to cytoskeleton organization and microtubule-based processes (Figure 6c). Enriched proteins are detected in different cellular compartments for upregulated and downregulated genes, respectively, such as the ER for upregulated genes, and the cytoskeleton for downregulated genes (Figure 6d). In terms of molecular function, RNA-binding proteins are highly enriched among upregulated genes, while actin-binding proteins are highly enriched among downregulated genes (Figure 6e).

**Figure 6.**
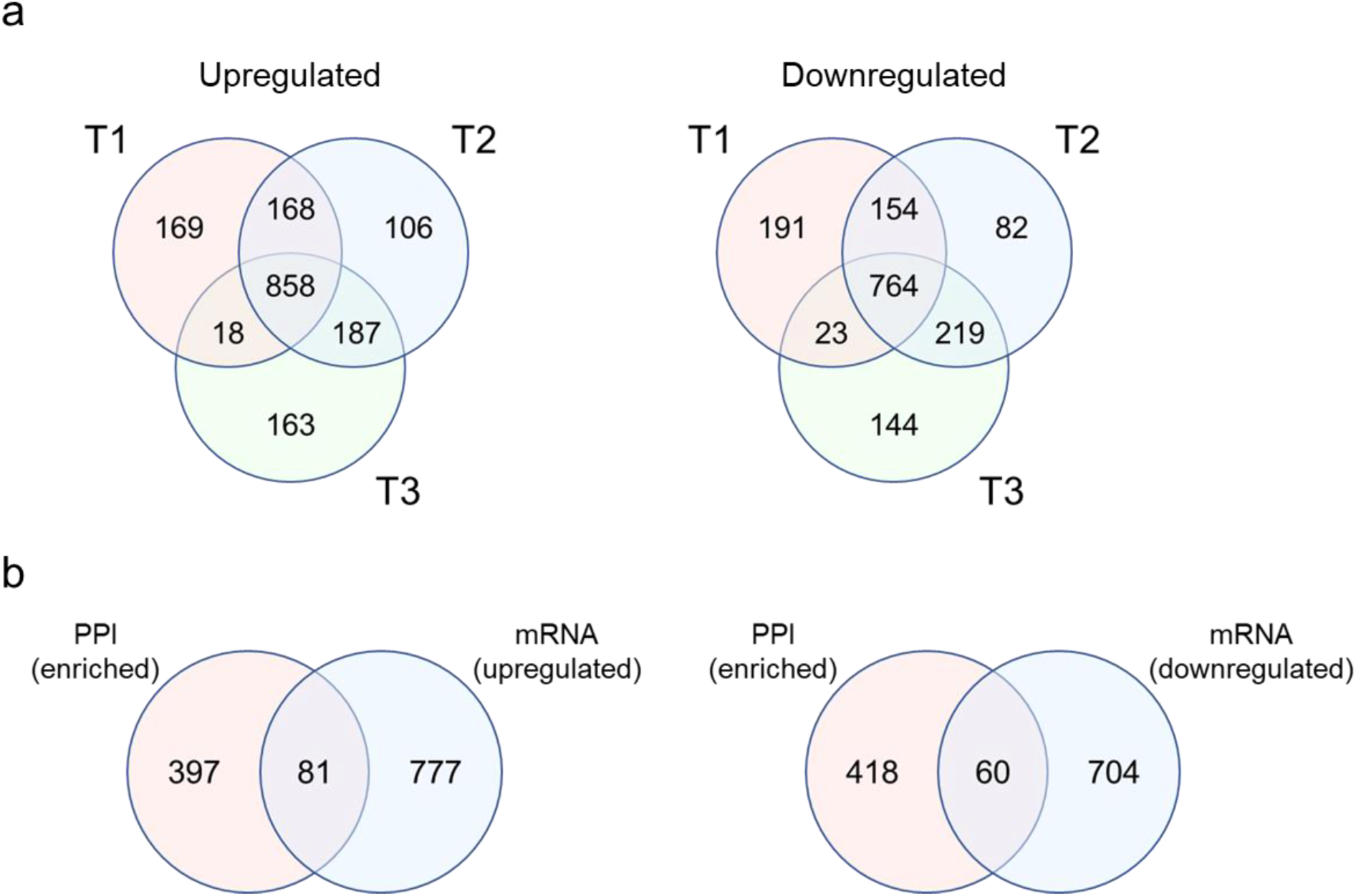
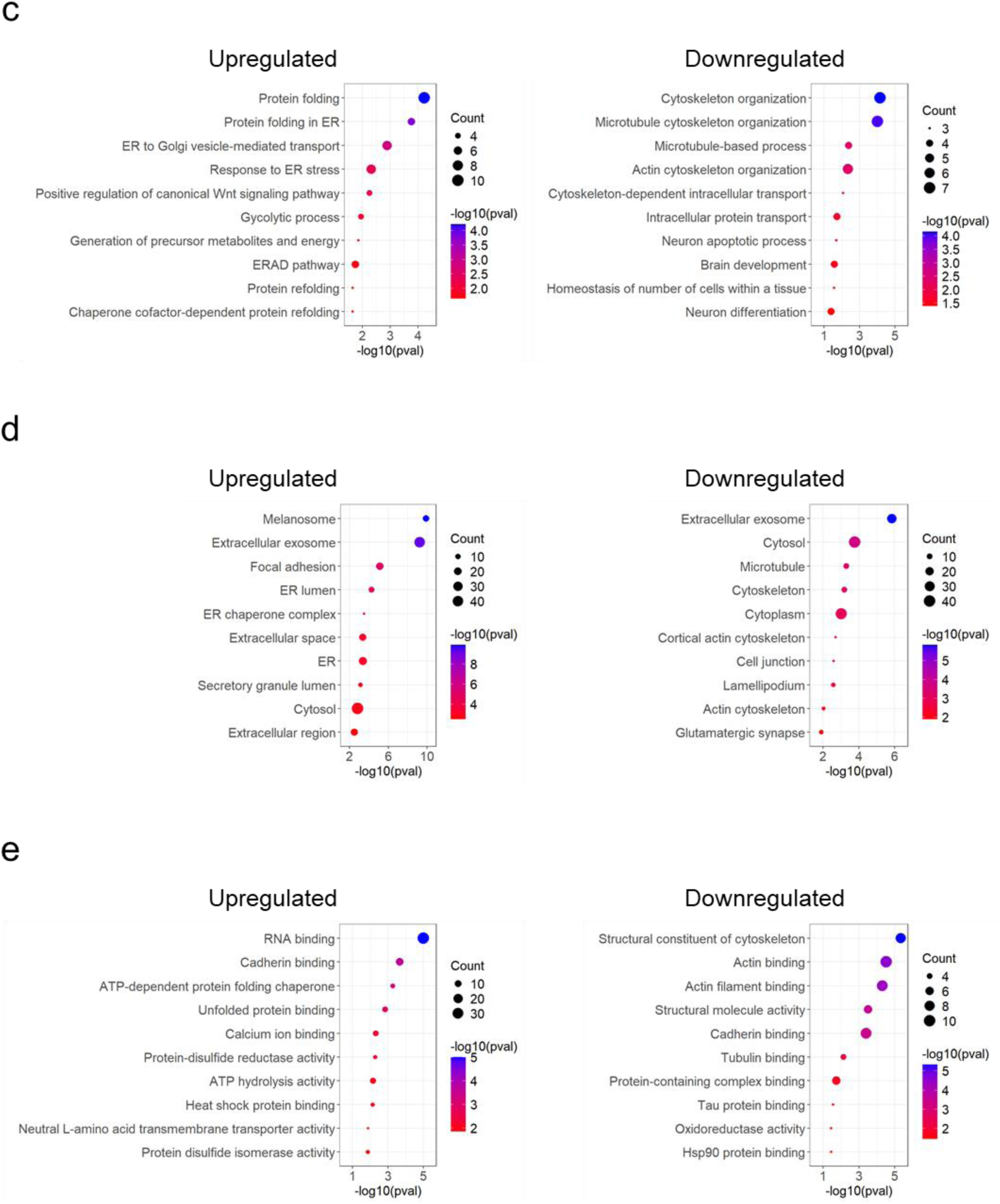
Transcriptional profiling and enriched protein-protein interactions following cell stimulation over time. (a) Numbers of upregulated and downregulated genes at each time point post-stimulation (T1, T2, and T3). (b) Numbers of differentially expressed genes encoding proteins enriched for IL-2 interactions (n=478) identified by proximity-based labeling. Gene ontology (GO) analysis of these differentially expressed, PPI-enriched genes categorized by (c) biological process, (d) cellular component, and (e) molecular function. All upregulated (n=1669) and downregulated (n=1577) genes identified by RNA sequencing were used as the background set. The top 10 GO terms were ranked by p-value.

To integrate RNA expression and PPI data in a time-resolved manner, we generated gene clusters based on expression changes from 0 to 8 hours. The maSigPro tool was used to identify genes with statistically significant temporal expression patterns [57]. Based on the changes of mRNA expression or enriched interaction with IL-2 proteins between each time point, a total of 27 and 23 clusters were identified in RNA sequencing and PPI data, respectively (p-value < 0.05) (Figure S7). A total of 57 proteins showed increased interaction with IL-2 from T0 to T1, enriching biological processes such as exosomal secretion and multivesicular body formation (Figure 7a). Between T1 and T2, 105 proteins exhibited enhanced interaction with IL-2, associated with processes including cytoskeleton organization and protein folding (Figure 7b). From T2 to T3, 29 proteins displayed increased interaction with IL-2, contributing to enrichment in cytoplasmic translation (Figure 7c). At time point T1, a total of 20 genes were identified as upregulated and enriched in IL-2 protein interactions, including CANX, HSPA5, HSP90B1, PDIA4, ERP44, and PTPN1 (Figure 7d), involved in protein folding in ER and response to ER stress. A total of 34 genes, including CANX, PDIA3, HSP90B1, HSP90AB1, HSPD1, HSPA8, HSPA9, and VCP, showed upregulated mRNA expression and enriched PPI at the same time point T2, resulting in enriched biological processes, such as protein folding in ER and response to unfolded protein (Figure 7e). Only 7 genes were detected at T3, including VCP gene, which is involved in ERAD (Figure 7f). None of the protein interactions were consistently enriched across all time points (Figure 7g). At both T1 and T2, protein folding in the ER was the most significantly enriched biological process in the GO analysis.

**Figure 7.**
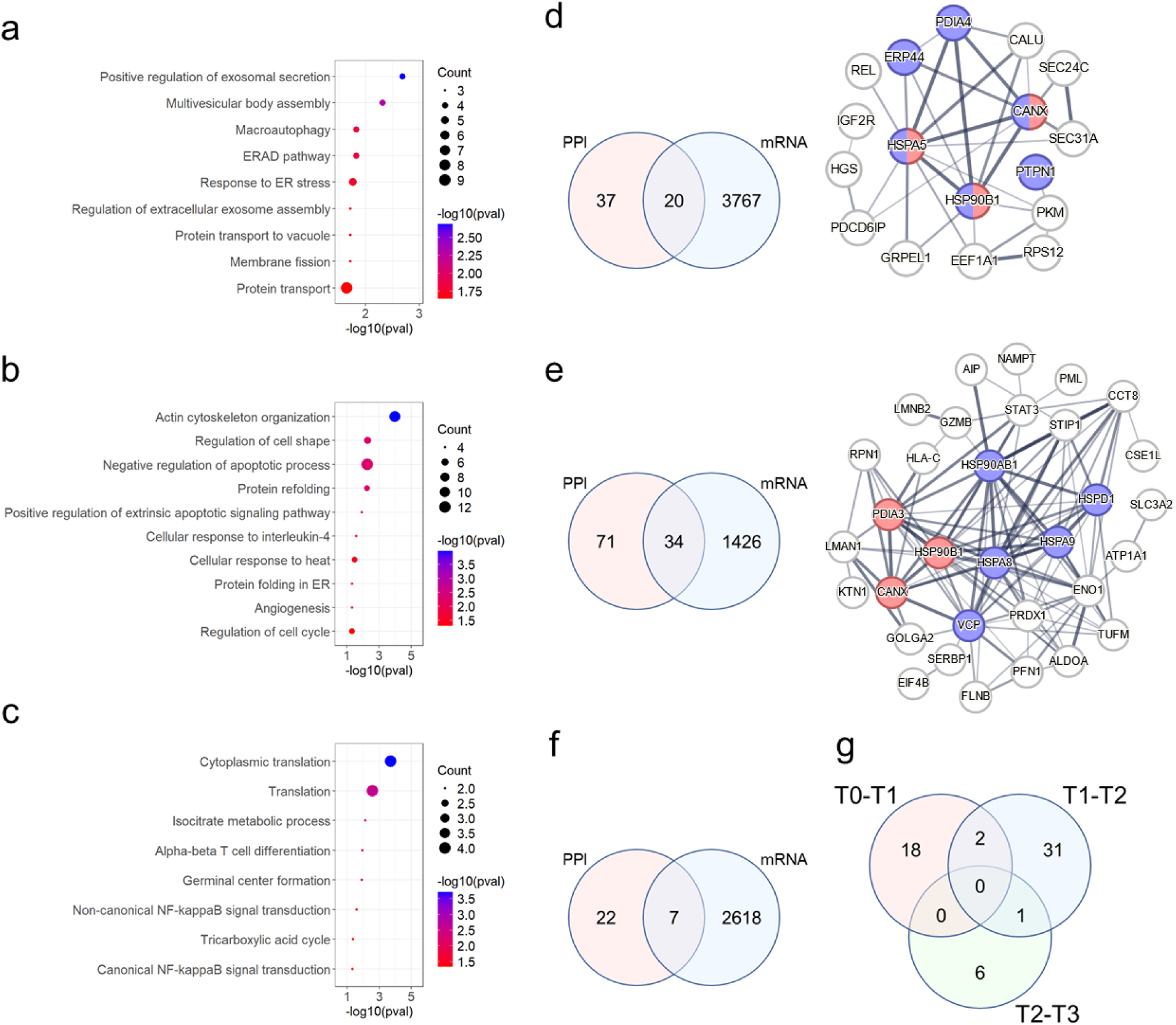
Temporal changes in transcriptional profiles and protein-protein interactions (PPIs). Gene ontology overrepresentation analysis of genes enriched in PPIs at each time interval: (a) T0–T1 (0–4 hours post-stimulation), (b) T1–T2 (4–6 hours post-stimulation), and (c) T2–T3 (6–8 hours post-stimulation). A total of 685 proteins showing significant changes in PPI enrichment over the time course were used as the background set. (d–f) Genes exhibiting both enriched PPIs and upregulated mRNA expression during: (d) T0–T1, (e) T1–T2, and (f) T2–T3. The STRING database was used to visualize significantly enriched protein-protein interactions. In T0–T1, red nodes indicate proteins involved in ER protein folding, and purple nodes indicate proteins involved in the ER stress response. In T1–T2, red nodes represent ER protein folding, and purple nodes represent the unfolded protein response. (g) Numbers of genes with upregulated mRNA expression and enriched PPI at each time interval post-stimulation (T0-T1, T1-T2, and T2-T3).

## Discussion

Recent advances in cytokine secretion profiling have enabled time-resolved and single-cell analyses through innovative labeling strategies, including oligo-barcoded antibody capture for sequencing-based detection and fluorescence-tagged antibody immobilization for real-time live-cell imaging [58,59]. Compared to direct labeling methods, which exploit specific binding interactions between molecules, proximity-based labeling methods can construct the physical network of proteins by detecting proteins within a specific range of target molecules. This feature makes it an optimal method for the profiling of interactions in protein secretion, which is mediated by a complex repertoire of secretory machinery proteins. We used the BAR method to capture the biotinylated proteins near the target protein, human cytokine IL-2, and quantified the PPIs by MS analysis. Enriched PPIs were detected in stimulated T cells, indicating that proteins that interact with IL-2 are successfully identified during the secretion of cytokine. We found that PPIs involved in conventional secretory pathways, comprising ER to Golgi transport, protein folding, and vesicle-mediated transport, are highly enriched in MS analysis (Figure 3a). In previous studies, IL-2 has been known to be secreted using a canonical secretory pathway from the ER, across the Golgi, and to the plasma cell membrane. However, only a few proteins, such as Rab3d, Rab19, Rab37, Syn6, and Vti1b, have been proven to interact and co-localize with IL-2 [46]. Therefore, this study provides a wealth of information on the systematic network of protein interactions in specific cytokine secretions, which can be exploited to develop therapeutic applications in diseases of the immune system.

Among the enriched secretory pathways, we found that protein folding in ER has the strongest association with the level of IL-2 secretion, validated by siRNA-mediated knockdown. Protein folding in the ER governs the secretion efficiency, functionality, and subcellular localization of both endogenous and recombinant proteins [60]. This early secretory pathway decides the fate of nascent proteins either being transported to Golgi with proper folding and assembling or inducing the unfolded protein response (UPR) and disposed by ERAD [61]. A plethora of proteins are involved in this process, including chaperones, lectins, redox enzymes, protein isomerases, and glycosylation enzymes, and the misfolded proteins generated from the malfunction of this system cause neurodegenerative diseases, such as Alzheimer’s and Parkinson’s [62]. In validation of target proteins related to the protein folding in ER, 10 out of 12 putative targets showed a significant reduction of IL-2 secretion when their expression was inhibited by siRNA (Figure 4c). Among the 10 PPIs enriched in IL-2 secreting T cells, HSPA8 and UGGT1 exhibited a reduction rate of more than 50% compared to the control (Figure 4c). HSPA8 is a molecular chaperone involved in various biological processes, such as proteostasis, autophagy, clathrin uncoating, and viral replication, and expressed constitutively without cellular stress unlike other chaperones [63]. Notably, HSPA8 proteins co-localize with IL-2 upon stimulation (Figure S3), and knockdown of the HSPA8 gene resulted in approximately a 70% reduction in IL-2 secretion (Figure 4c), suggesting a potential role for HSPA8 in regulating IL-2 secretion that requires further investigation.

The expression of cytokines is tightly regulated in a time-dependent manner within a few hours after immune cell stimulation [64]. This multilayered regulation involves transcriptional, post-transcriptional, and post-translational control mediated by transcription factors, RNA-binding proteins, and other secretory machinery proteins. Therefore, systematic approaches are essential to understand cytokine secretion as a whole process governed by the physical and functional network of PPIs. RNA-binding proteins, which are not directly related to protein secretion, are detected at different time points after stimulation (Figure 5c). This indicates that the HRP-mediated labeling method can detect co-localization of IL-2 proteins and IL-2 mRNA-binding proteins, given the labeling range of this method (200-300 nm) [14]. Among 15 RNA-binding proteins, 9 of them (CAPRIN1, ATXN2L, CSDE1, CORO1A, TUFM, ANXA11, UBAP2L, ZC3HAV1, and STRAP) were previously identified showing strong interaction with IL-2 3’ UTR of in vitro transcribed RNA in pull-down assay [65]. Knockdown of ATXN2L or ANXA11 significantly decreased IL-2 secretion at both 8 h and 24 h after stimulation (Figure 5d), confirming the function of those RNA-binding proteins in IL-2 production.

The stimulation of human T cells induces the changes of transcriptional profile, thereby regulating immune response, such as cytokine secretion [66]. We found that biological pathways such as protein transport from the ER to the Golgi and protein folding in the ER were significantly enriched in PPIs alongside upregulated gene expression (Figure 6c). This indicates that cell stimulation enhances the expression of genes involved in protein transport and folding. In contrast, cytoskeleton organization and microtubule-based processes were enriched in PPIs together with downregulated gene expression (Figure 6c), suggesting that stimulation suppresses genes involved in cytoskeletal regulation. Interestingly, although cytoskeleton-related genes were downregulated after T cell stimulation, the corresponding proteins showed increased enrichment within IL-2 interaction networks. We also employed clustering based on the changes of protein interaction and RNA expression between specific time points (0-4, 4-6, 6-8 h after stimulation). As a result, 20, 34, and 7 protein interactions were enriched at only T1, T2, and T3, respectively (Figure 7d-f). It is worth noting that there is minimal overlap in protein interactions between each pair of time points, two between T1 and T2, one between T2 and T3, and none between T3 and T1, suggesting highly dynamic and spatiotemporally distinct interactions between these proteins and IL-2 (Figure 7g).

The BAR method has proven effective for identifying PPIs within native cellular environments, particularly at the nuclear envelope [40]. It has also been applied to uncover interaction networks of secreted endogenous and recombinant proteins, thereby identifying potential engineering targets for modulating protein secretion [67–69]. However, it has several limitations. First, it features a relatively large labeling radius, which can result in indirect or non-specific labeling of proteins that are merely proximal rather than true interactors [70]. Second, the biotinylation reaction is spatially restricted, primarily targeting extracellular proteins or those within the secretory pathway and therefore cannot comprehensively capture intracellular interactions. This constraint stems from the use of HRP, which catalyzes radical formation in the presence of hydrogen peroxide (H₂O₂) to label nearby electron-rich amino acids. HRP functions only in oxidizing environments, such as the ER or Golgi lumen and the extracellular space, because its structural integrity depends on disulfide bonds and calcium-binding sites, which are disrupted under the cytosol’s reducing conditions [71]. Consequently, HRP-based proximity labeling is unsuitable for detecting protein interactions within the cytoplasm or other reducing compartments. Furthermore, reliance on biotinylation can introduce bias, as labeling efficiency depends on the accessibility and abundance of lysine residues on target proteins, potentially leading to uneven or incomplete labeling across the proteome. To overcome these limitations, BAR can be complemented with affinity purification methods, such as pull-down assays, which selectively enrich interacting partners [72]. Moreover, alternative proximity labeling approaches with varied labeling radii, such as APEX and BioID, can be employed to achieve broader or more compartment-specific interactome coverage. Recently, a multiscale labeling strategy employing three distinct chemical reactions was introduced to map protein interactions with varying spatial resolutions and labeling ranges [73].

## Conclusion

In this study, we demonstrated the utility of the BAR method for capturing a comprehensive network of PPIs surrounding IL-2 during its secretion in human T cells. By integrating proximity-based labeling with mass spectrometry, we identified dynamic and stage-specific interactomes that include not only canonical secretory machinery but also regulatory proteins, such as chaperones and RNA-binding proteins. Our findings highlight the critical role of ER-associated protein folding in IL-2 secretion, with functional validation supporting the importance of specific chaperones like HSPA8. Furthermore, the detection of RNA-binding proteins with known affinity to the IL-2 3′ UTR suggests that BAR is capable of capturing spatial co-localization events between cytokine proteins and their mRNA-binding partners. Temporal clustering analysis revealed that IL-2 interacts with distinct sets of proteins at different stages of secretion, reflecting tightly regulated spatial and temporal dynamics. Although the BAR method presents inherent limitations, such as a relatively broad labeling radius, spatial restrictions to oxidizing environments, and biotinylation biases, its integration with complementary techniques and advanced labeling strategies offers a powerful platform for dissecting complex secretory processes. Collectively, our results provide a systems-level view of cytokine secretion and underscore the value of proximity labeling in elucidating the regulatory landscape of immune responses, with potential implications for therapeutic targeting of immune-related diseases.

## Acknowledgments

This work was supported by generous funding from NIH (R35 GM119850), NSF (CBET-2030039), and the Novo Nordisk Foundation (NNF20SA0066621). Fluorescence microscopy was performed at UCSD School of Medicine Microscopy Core, which is supported by Grant P30 (NINDS P30NS047101). Mass spectrometry analysis was performed by the Sanford Burnham Prebys Proteomics Core, with assistance from Dr. Svetlana Maurya. This work also includes data generated at the UC San Diego IGM Genomics Center utilizing an Illumina NovaSeq X Plus that was purchased with funding from a National Institutes of Health SIG grant (#S10 OD026929).

## Data Availability

The study findings are supported by data available upon request from the corresponding author. Raw and processed mass spectrometry proteomics data have been deposited in the MassIVE repository (ProteomeXchange Consortium; https://doi.org/10.25345/C5TT4G61Q; reference PXD068039). Raw and processed transcriptomic data are available in GEO under the dataset identifier GSE306928.

