## Supporting Information for "Systematic Mapping of Protein Interactions Underlying IL-2 Secretion in Human T Cells"

**Supplementary Figures**

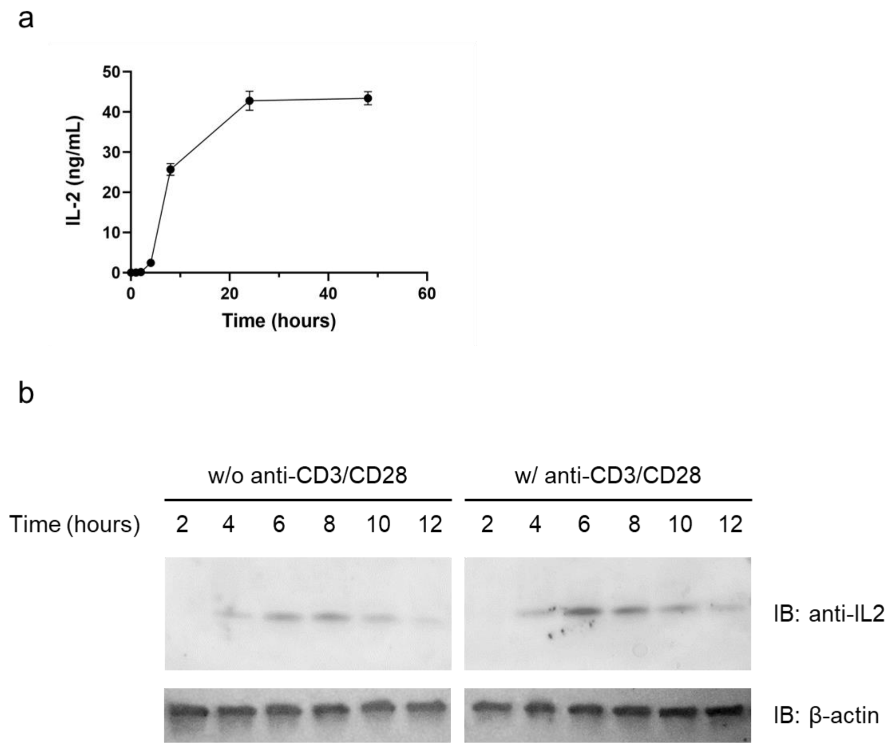


**Supplementary Figure 1.** IL-2 expression following stimulation of Jurkat T cells. (a) Concentration of secreted IL-2 in the culture supernatant measured by ELISA after stimulation. (b) Intracellular IL-2 protein levels at various time points post-stimulation, measured by Western blot in the presence or absence of anti-CD3/CD28 antibodies.

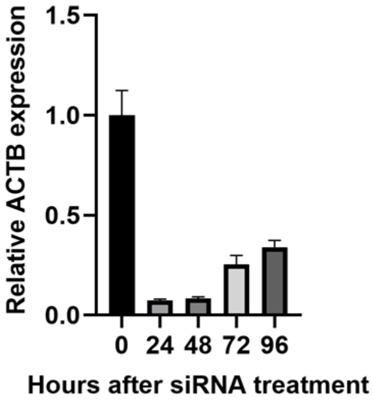


**Supplementary Figure 2.** Relative ACTB gene expression levels at different time points after siRNA treatment.

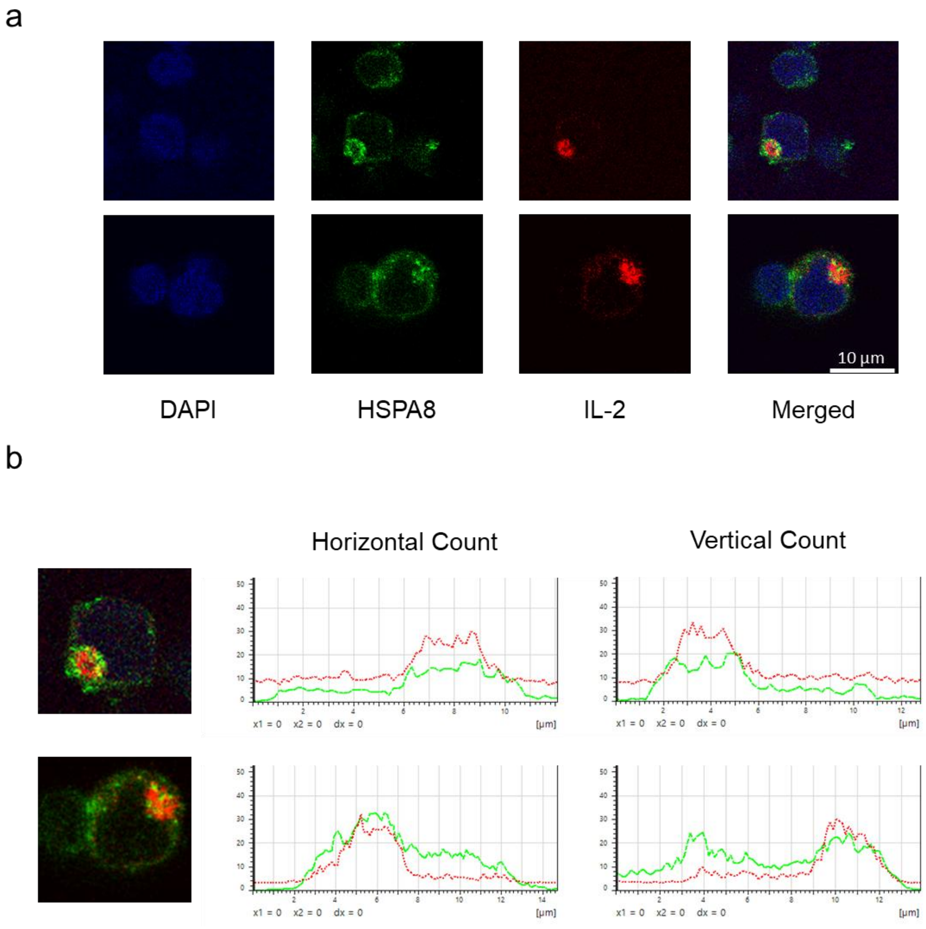


**Supplementary Figure 3.** Protein–protein interaction between IL-2 and HSPA8. (a) Co-localization of IL-2 and HSPA8 proteins in stimulated Jurkat T cells. (b) Line profile analysis showing horizontal and vertical signal intensity of HSPA8 and IL-2. Intracellular HSPA8 was visualized using DyLight 594 (green), and IL-2 was stained with DyLight 650 (red).

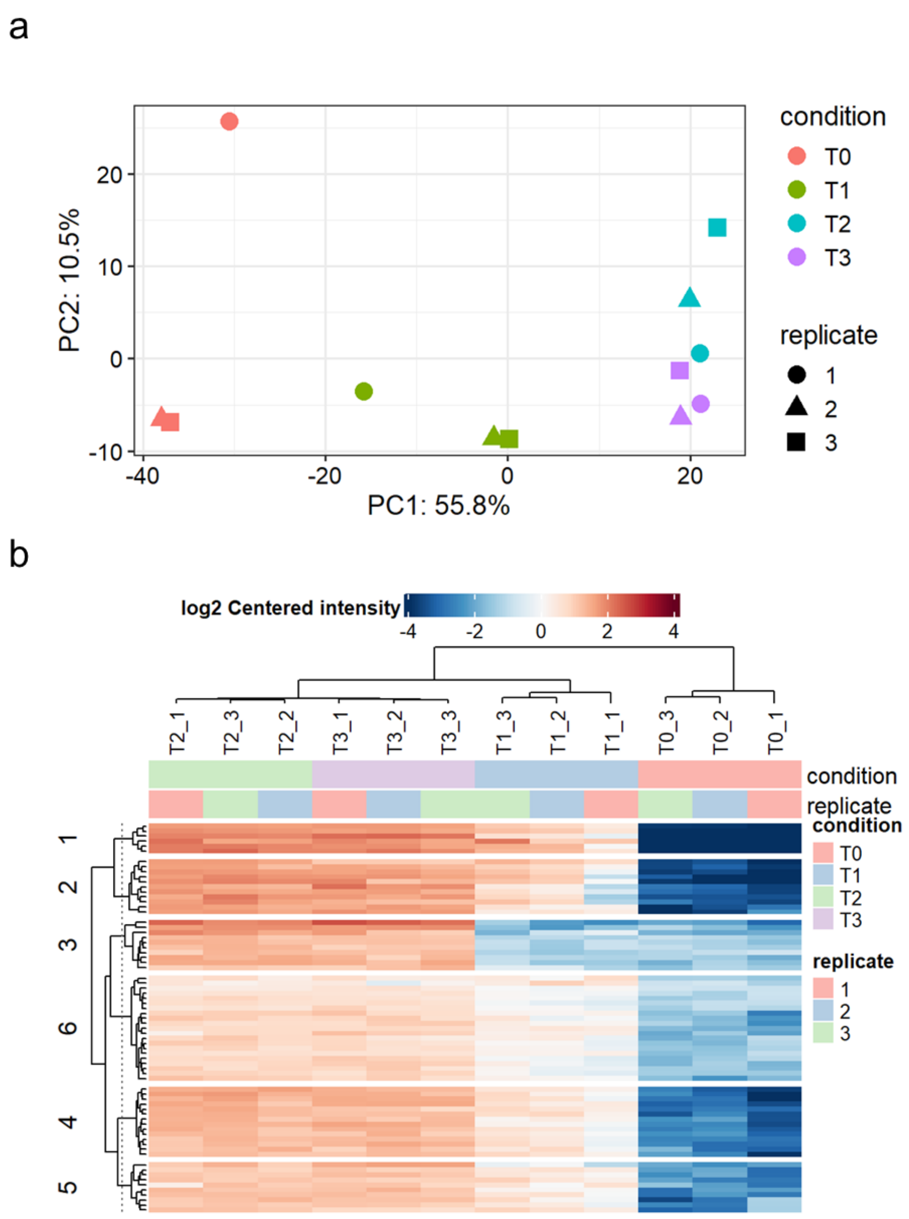


**Supplementary Figure 4.** Correlation of samples at different time points identified by proximity-based labeling. (a) Principal component analysis (PCA) of 12 samples (triplicates for each time point): T0 (control), T1 (4 hours post-stimulation), T2 (6 hours post-stimulation), and T3 (8 hours post-stimulation). (b) Heatmap showing sample clustering across the same time points.

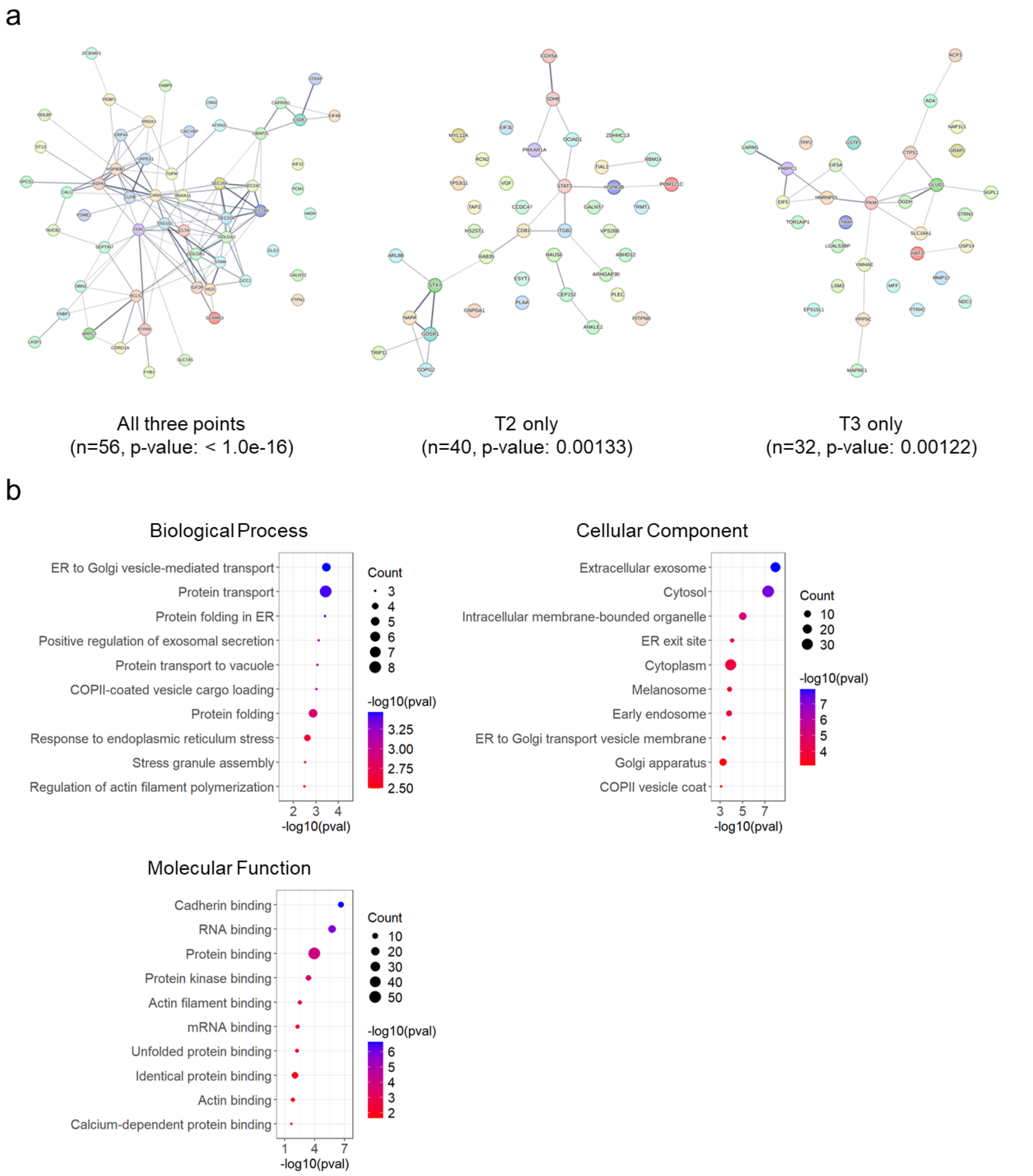


**Supplementary Figure 5.** Enriched protein–protein interactions and biological pathways across time points. (a) STRING network analysis of enriched proteins identified at all three time points (T1, T2, and T3), uniquely at T2, or uniquely at T3. (b) Gene ontology analysis of 56 proteins consistently enriched at all three time points (T1, T2, and T3).

a

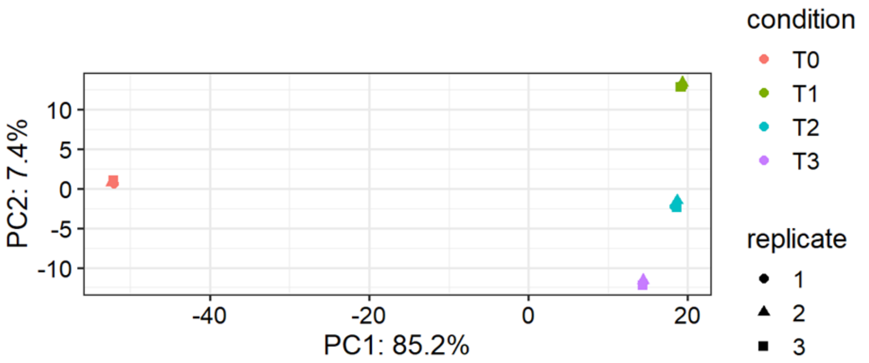


b

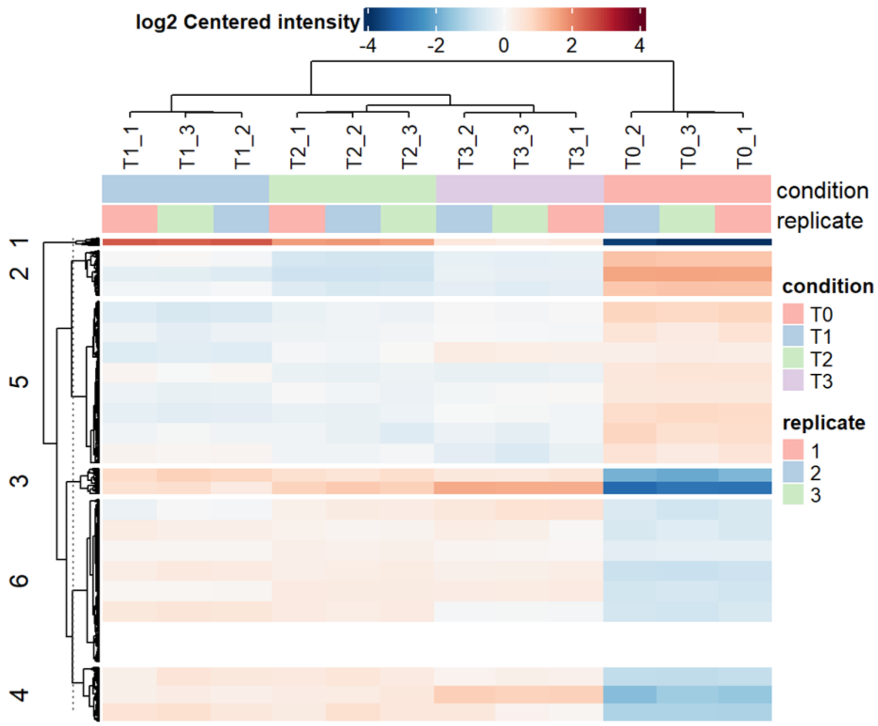


**Supplementary Figure 6.** Correlation of samples across different time points based on RNA sequencing. (a) Principal component analysis (PCA) of 12 samples (triplicates per time point): T0 (control), T1 (4 hours post-stimulation), T2 (6 hours post-stimulation), and T3 (8 hours post-stimulation). (b) Heatmap showing hierarchical clustering of the same samples, highlighting transcriptomic similarity among T0, T1, T2, and T3.

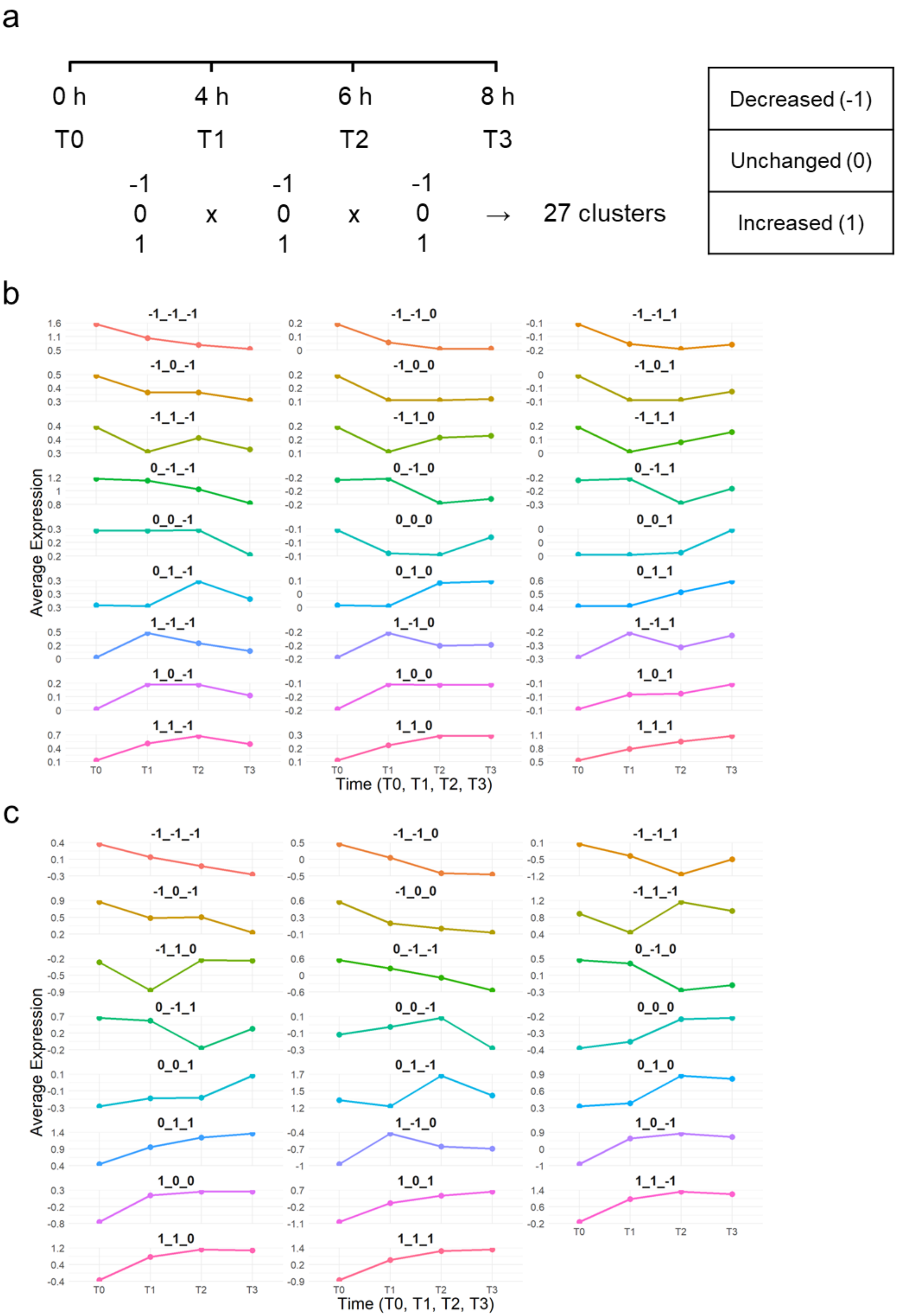
a
0 h
4 h
6 h
8 h
T0
T1
T2
T3
-1 0 1 x -1 0 1 x -1 0 1 → 27 clusters
Decreased (-1)
Unchanged (0)
Increased (1)
b
-1_-1_-1
-1_-1_0
-1_-1_1
-1_0_-1
-1_0_0
-1_0_1
-1_1_-1
-1_1_0
-1_1_1
0_-1_-1
0_-1_0
0_-1_1
0_0_-1
0_0_0
0_0_1
0_1_-1
0_1_0
0_1_1
1_-1_-1
1_-1_0
1_-1_1
1_0_-1
1_0_0
1_0_1
1_1_-1
1_1_0
1_1_1
Average Expression
Time (T0, T1, T2, T3)
c
-1_-1_-1
-1_-1_0
-1_-1_1
-1_0_-1
-1_0_0
-1_1_-1
-1_1_0
0_-1_-1
0_-1_0
0_-1_1
0_0_-1
0_0_0
0_0_1
0_1_-1
0_1_0
0_1_1
1_-1_0
1_0_-1
1_0_0
1_0_1
1_1_-1
1_1_0
1_1_1
Average Expression
Time (T0, T1, T2, T3)

**Supplementary Figure 7.** Generation of gene clusters based on significant changes between time points. (a) A total of 27 distinct clusters were defined by combining three differential expression/enrichment states(decreased, unchanged, and increased) across three consecutive time intervals (T0–T1, T1–T2, and T2–T3). (b) Expression patterns of mRNA across the defined clusters. (c) Enrichment trends of protein-protein interactions corresponding to each cluster.
